# Inhibition of USP4 attenuates epithelial-mesenchymal transition of renal tubular epithelial cells by TβRI

**DOI:** 10.1101/2020.10.28.358796

**Authors:** Jin-yun Pu, Yu Zhang, Li-xia Wang, Jie Wang, Jian-hua Zhou

## Abstract

The process of epithelial-mesenchymal transition (EMT) is required for the progression of renal interstitial fibrosis (RIF). Ubiquitin-specific protease 4 (USP4) can facilitate development of transforming growth factor, beta 1 (TGF-β1) induced EMT in some cancer cells. However, the role of USP4 in EMT during RIF remains unknown. We aimed to explore the effect of USP4 on the EMT induced by TGF-β1 of renal tubular epithelial cells and involved mechanism in RIF. In vivo, on the 7th and 14th day after unilateral ureteral obstruction (UUO), the expression of USP4 protein in the obstructed kidneys was detected by immunohistochemistry and Western blot assay. In vitro, NRK-52E cells were stimulated with TGF-β1 10ng/ml. The protein expressions of USP4, E-cadherin and alpha smooth muscle actin (α-SMA) were detected at different time points by Western blot. After transfected with USP4 siRNA, the cells were cultured with TGF-β1 for additional 24 hours. The expressions of E-cadherin, α-SMA, and TGFβ receptor type I (TβRI) were detected by immunofluorescence. And the protein expressions of USP4, E-cadherin, α-SMA and TβRI were detected by Western blot assay. Compared with sham operation group, the expression of USP4 in UUO model group increased significantly with the prolongation of obstruction time. After NRK-52E was stimulated by TGF-β1, the expression of USP4 protein increased gradually. At 6h, 12h, and 24h, the difference between the experimental group and the control group was statistically significant. At the same time, E-cadherin decreased significantly, while α-SMA increased significantly. Compared with the TGF-β1 group, the cells in USP4 siRNA transfection group restored E-cadherin and weakened α-SMA expression. At the same time, protein expressions of USP4 and TβRI were also significantly decreased. These data imply that USP4 is a harmful molecule induced by TGF-β1, which plays an important role by upregulating the expression of TβRI and promoting EMT of renal tubular epithelial cells, thereby facilitating renal interstitial fibrosis.

## 1. Introduction

Renal interstitial fibrosis (RIF) is the final common pathological pathway of almost all chronic kidney diseases, which is closely related to poor clinical prognosis[1]. RIF is characterized by the deposition of excessive extracellular matrix and accumulation of myofibroblasts in the interstitial area[2]. Although great efforts have been made to find the molecular and cellular mechanisms underlying the development of RIF, there is still no effective treatment to prevent the occurrence and development of RIF.

Tubular epithelial cells are the main components of renal parenchyma and are often the target of various injuries[3]. Many studies have observed the phenomenon of epithelial-mesenchymal transition (EMT) during RIF[4], which contributes to the myofibroblasts pool. EMT refers to the process that tubular epithelial cells lose epithelial markers E-cadherin and obtain mesenchymal markers, such as fibronectin, vimentin, and alpha smooth muscle actin (α-SMA). Among them, α-SMA is an accepted marker for EMT and fibrosis[5]. Shreds of evidence show that inhibiting EMT can alleviate RIF[6].

Ubiquitination refers to the process of binding ubiquitin covalently to target proteins under the catalysis of multi-step cascade enzymes, which is one of the critical post-translational modifications. Different from the single group added by phosphorylation, methylation and acetylation, ubiquitin is a small protein composed of 76 amino acids, which widely exists in all eukaryotic cells. Ubiquitination is not only involved in the regulation of protein quantity but also plays an important role in the regulation of protein activity, protein-protein interaction, and protein subcellular localization[7]. Ubiquitination is widely involved in various cellular processes, including cell proliferation, apoptosis, autophagy, endocytosis, and immune response[8]. The regulation of ubiquitination pathway has been considered as a promising therapeutic strategy for many diseases[9]. Ubiquitination is also strictly reversibly controlled. Deubiquitination is a process of protein hydrolysis in which ubiquitin is removed directly from protein substrates by deubiquitinases (DUBs). There are many DUBs in many cells. According to the sequence and structure, DUBs are divided into five families[10]. Ubiquitin-specific proteases (USPs), the largest subfamily of DUBs, account for more than 50% of DUBs and there are more than 50 members[11]. DUBs can not only inhibit the ubiquitination process but also promote the ubiquitination process by decomposing ubiquitination inhibitors, recycling ubiquitin molecules, and proofreading the ubiquitination process, thus forming a complex network covering almost all cell function together with ubiquitination system[12].

Ubiquitin-specific protease 4 (USP4), also known as UNP or Unph, is a member of the USPs family, which is the first DUBs identified in mammalian cells[13]. USP4 is an important DUB, which can inhibit its degradation or change its characteristics by recognizing specific target proteins and deubiquitinating them. It plays an important regulatory role in tumors, virus infection, and a variety of signaling pathways[13]. Previous studies have found that this enzyme acts as an oncoprotein or tumor suppressor in cancer biology[14]. Peter et al. discover that USP4 can promote the EMT of breast cancer cells induced by transforming growth factor, beta 1 (TGF-β1)[15]. However, The role of USP4 in epithelial-mesenchymal transition in renal interstitial fibrosis has not been reported yet.

In the present study, we aimed to investigate that if USP4 exerts an effect on TGF-β1 induced EMT in tubular epithelial cells during renal interstitial fibrosis.

## 2. Method

### 2.1 Unilateral ureteral obstruction model

Twenty male Wistar rats weighing 200 to 220g were obtained from Beijing Vital River Experimental Animals Technology Ltd. (Beijing, China) and housed under SPF-laboratory conditions. Unilateral ureteral obstruction (UUO) was established following an approved protocol by the Institutional Animal Care Committee and Use Committee of Tongji Hospital, Tongji Medical College, Huazhong University of Science and Technology, as described previously[16]. In brief, the upper and lower third of the left ureter were ligatured with 4-0 silk sutures, and then the middle third was cut off. As controls, sham-operated rats underwent the same surgery except for ureter ligation and division. To investigate the severity and time course of UUO-induced renal fibrosis, five rats in sham and UUO groups were sacrificed separately on the 7th or 14th day after the operation. And then, the left kidneys were harvested. Formalin-fixed paraffin-embedded kidney tissues were prepared for histological evaluation. Other parts were frozen immediately in liquid nitrogen and stored for protein extraction.

### 2.2 Masson’s trichrome staining and immunohistochemistry

The acquired kidney was prepared as formalin-fixed paraffin-embedded tissue. Then the tissue samples were sliced to 10μm thickness and then dewaxed. Masson’s trichrome staining was performed on slices to visualize fibers according to standard protocol. After dehydrated, the slices were boiled for 20 minutes in citrate buffer for antigen retrieval. The non-specific antigen was blocked by 5% BSA for 60 minutes at room temperature. Then, the tissue sections were incubated with the primary antibody against USP4 (Cell Signaling Technology) diluted in 5% BSA at 4°C overnight. After that, the slices were washed in PBS three times. For immunohistochemistry, the HRP-conjugated secondary antibody was incubated for 60 minutes at room temperature and then performed with DAB assay according to the instruction.

### 2.3 Cell culture and treatment

Rat tubular epithelial cells (NRK-52E) obtained from the American Type Culture Collection were maintained in DMEM culture media/high glucose with 10% fetal bovine serum (Gibco) in a humidified incubator with 5% CO2 at 37°C. Cells were cultured without or with human TGF-β1 (PeproTech) at a concentration of 10 ng/ml for 15min, 30min, 1h, 3h, 6h, 12h, and 24h.

### 2.4 USP4 siRNA transfection

The 21-nucleotide small interfering RNA (5′-UAGACUGACCCAGCUGGAAdTdT-3′) targeting USP4 (USP4 siRNA) was designed and synthesized (Viewsolid). LipojetTM reagent was used to transfect USP4 siRNA molecules to NRK-52E cells according to kit instructions (SignaGen). Cells were plated the day before transfection so that the monolayer cell density reached 50% confluency. The lipoJetTM/USP4 siRNA transfection mix was prepared for 10 min at room temperature. Check siRNA silencing efficiency 6 hours post-transfection. Then the cells were stimulated with TGF-β1 for additional 24 hours.

### 2.5 Immunofluorescence

The NRK-52E cells were fixed in fresh 4% paraformaldehyde for 10 min in the culture plate at room temperature, washed with PBS for three times, permeabilized with 0.5% Triton X-100 for 5 min, and blocked with 1% bovine serum albumin for an hour. Then, the cells were incubated with primary antibodies at 4°C overnight in a dark humidity chamber. The cells were washed sufficiently with PBS and then incubated with secondary antibodies for an hour at room temperature in dark, followed by washing. Then the cells were stained with DAPI and mounted on glass slides and viewed with Olympus optical fluorescence microscope.

### 2.6 Western blot

Total protein from tissue or cells was extracted by RIPA lysis buffer. After denaturing, the samples were loaded in the Tris-gel for electrophoretic separation. After that, the protein was transferred onto the PVDF membrane. Then the membrane was incubated at 4°C overnight, with primary antibodies against E-cadherin (BD), α-SMA (Boster), TβRI (Proteintech), and GAPDH (CST). Secondary antibodies were incubated and ECL assay was performed. Relative amounts of each protein were compared in each group.

### 2.7 Statistical analysis

All experiments were carried out independently at least three times. All values were presented as mean ± standard deviation. The statistical analyses were performed using one-way analysis of variance (ANOVA) by SPSS 19.0 software, and p value <0.05 was considered significant statistically.

## 3. Results

### 3.1 USP4 expression increases in UUO rats

To examine the role of USP4 in renal fibrosis, dynamic expression of USP4 protein in established unilateral ureteral obstruction rats was analyzed. Masson’s trichrome staining showed a significant increase of collagen components in renal interstitium after obstruction for 7 or 14 days (Fig. 1A). Immunohistochemistry results showed that the expression of USP4 protein in UUO rats was significantly upregulated compared with the sham group as early as on day 7 and continued (Fig. 1B). Western blot assay was performed to quantify USP4 protein expression during the development of renal fibrosis. The results showed that USP4 expression on the 14th day was more than that on the 7th day. However, protein expression of USP4 in sham rats was rare (Fig. 1C). In this experiment, we found that expression of USP4 protein increased along with renal fibrosis in UUO rats.

**Figure 1.**
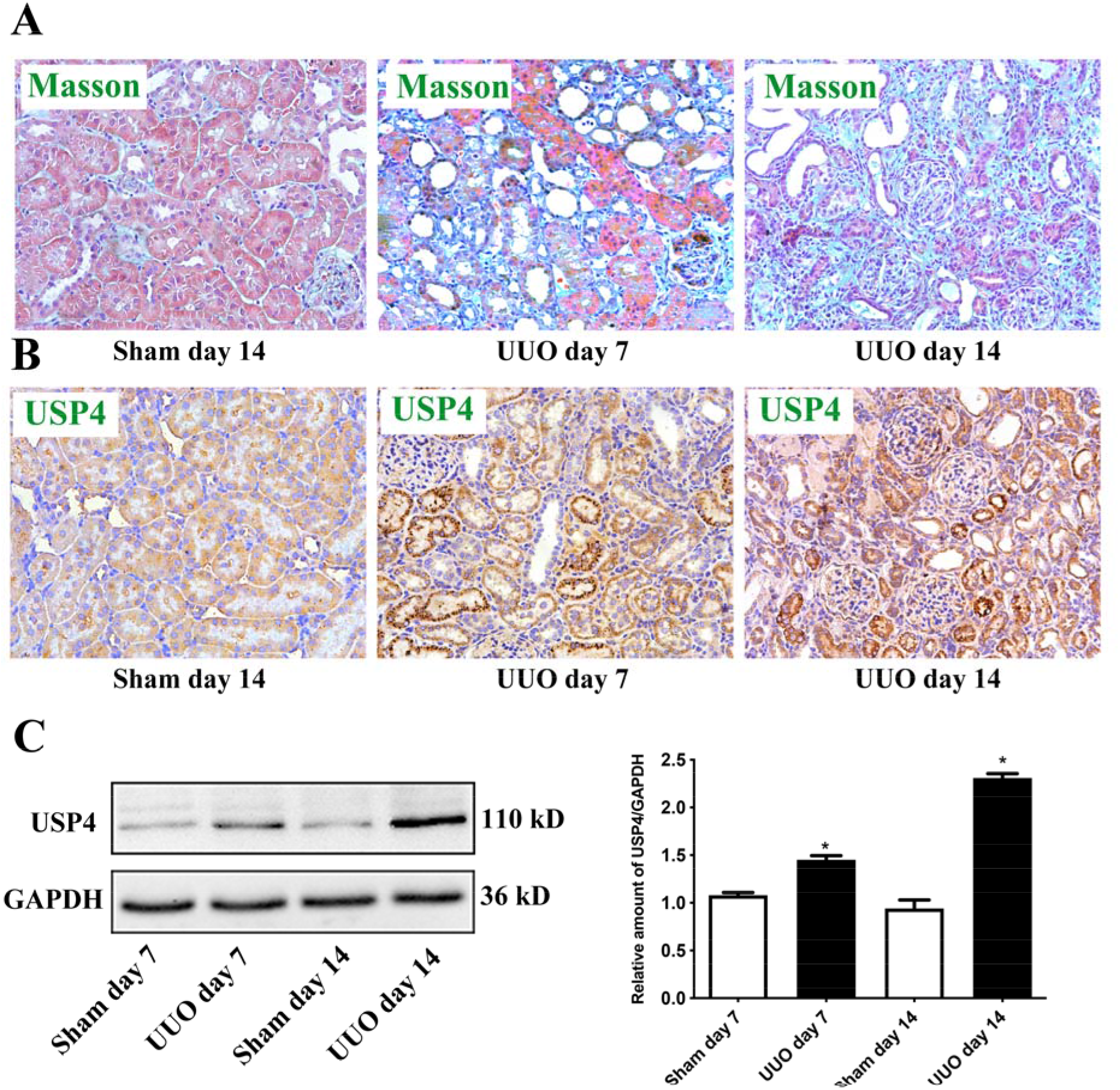
Expression of USP4 protein during renal fibrosis in sham and UUO rats. (A) Masson’s staining of renal fibrosis, magnification ×200. (B) Immunohistochemistry staining of USP4 expression, magnification ×200. (C) Western blot analysis of USP4 expression in the kidney of sham and UUO operated rats, the bars showed the relative expression quantity normalized to GAPDH and compared to sham rats. Representative data from five rats were shown as mean ± standard deviation. *p<0.05 compared to sham rats group.(null)

### 3.2 USP4 expression upregulates during EMT induced by TGF-β1 in NRK-52E cells

TGF-β1 is considered as one of the important inducers of EMT during renal fibrosis. In order to investigate the expression of USP4 on EMT in tubular epithelial cells, we performed in vitro experiment to investigate the expression level during EMT of NRK-52E cells. After treated with TGF-β1, E-cadherin was decreased, while α-SMA and USP4 were increased in NRK-52E cells (Fig. 2A). Quantitative analysis using western blot assay showed that USP4 was significantly upregulated after 6 hours in cells stimulated with TGF-β1 (Fig. 2B). This experiment indicated that NRK-52E cells upregulated expression of the USP4 protein during EMT.

**Fig. 2.**
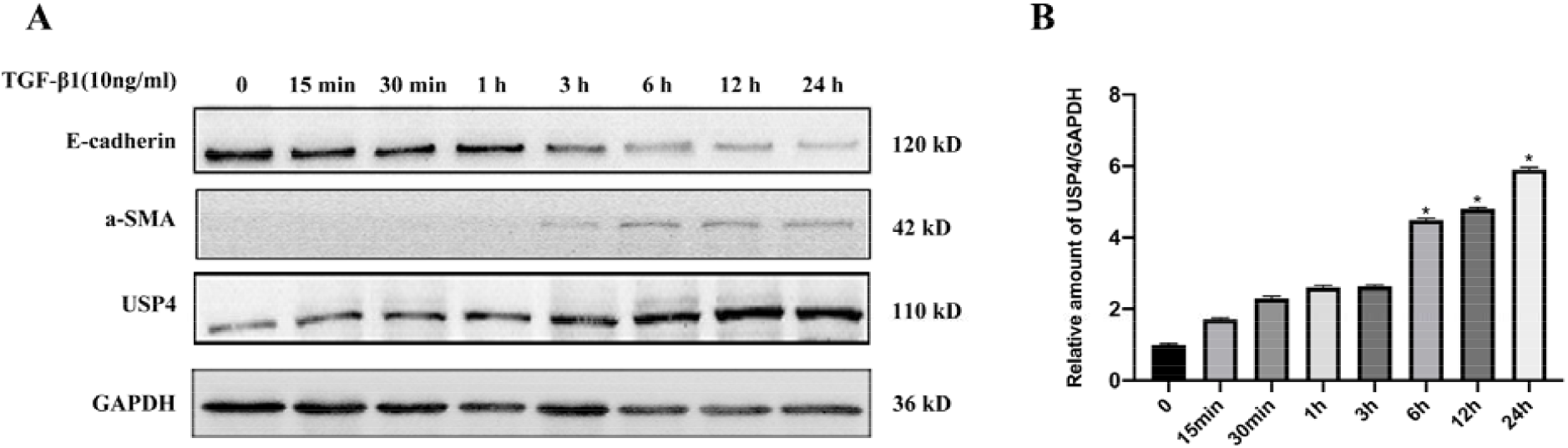
USP4 expression in TGF-β1-induced EMT in a time-dependent manner. NRK-52E cells were treated with TGF-β1 at concentrations of 10 ng/ml at extended periods (0, 15min, 30min, 1h, 3h, 12h, and 24h). (A) Protein expressions of E-cadherin, α-SMA, and USP4 by western blot assay in NRK-52E cells. (B) Western blot analysis of USP4 expression in NRK-52E cells at distinct time points. Bars showed relative expression quantity normalized to GAPDH compared to normal control cells without TGF-β1 treatment. *P<0.05 compared with normal control cells.

### 3.3 USP4 depletion inhibits TGF-β1-induced EMT in vitro

To investigate the effect of USP4 on EMT, we transfected USP4 siRNA to knockdown USP4 in NRK-52E cells and an immunofluorescence assay was applied to visualize the expression of E-cadherin and α-SMA in NRK-52E cells. After treatment with 10 ng/ml TGF-β1 for 24h, NRK-52E cells displayed downregulated E-cadherin but upregulated α-SMA expressions. Pretreatment with si-USP4, NRK-52E cells could restore E-cadherin while weaken α-SMA expression (Fig. 3A). Quantitative analysis using western blot assay showed that USP4 siRNA increased E-cadherin and decreased α-SMA significantly in TGF-β1 stimulated NRK-52E cells (Fig. 3B). These results indicated that si-USP4 inhibited EMT induced by TGF-β1 in NRK-52E cells.

**Fig. 3.**
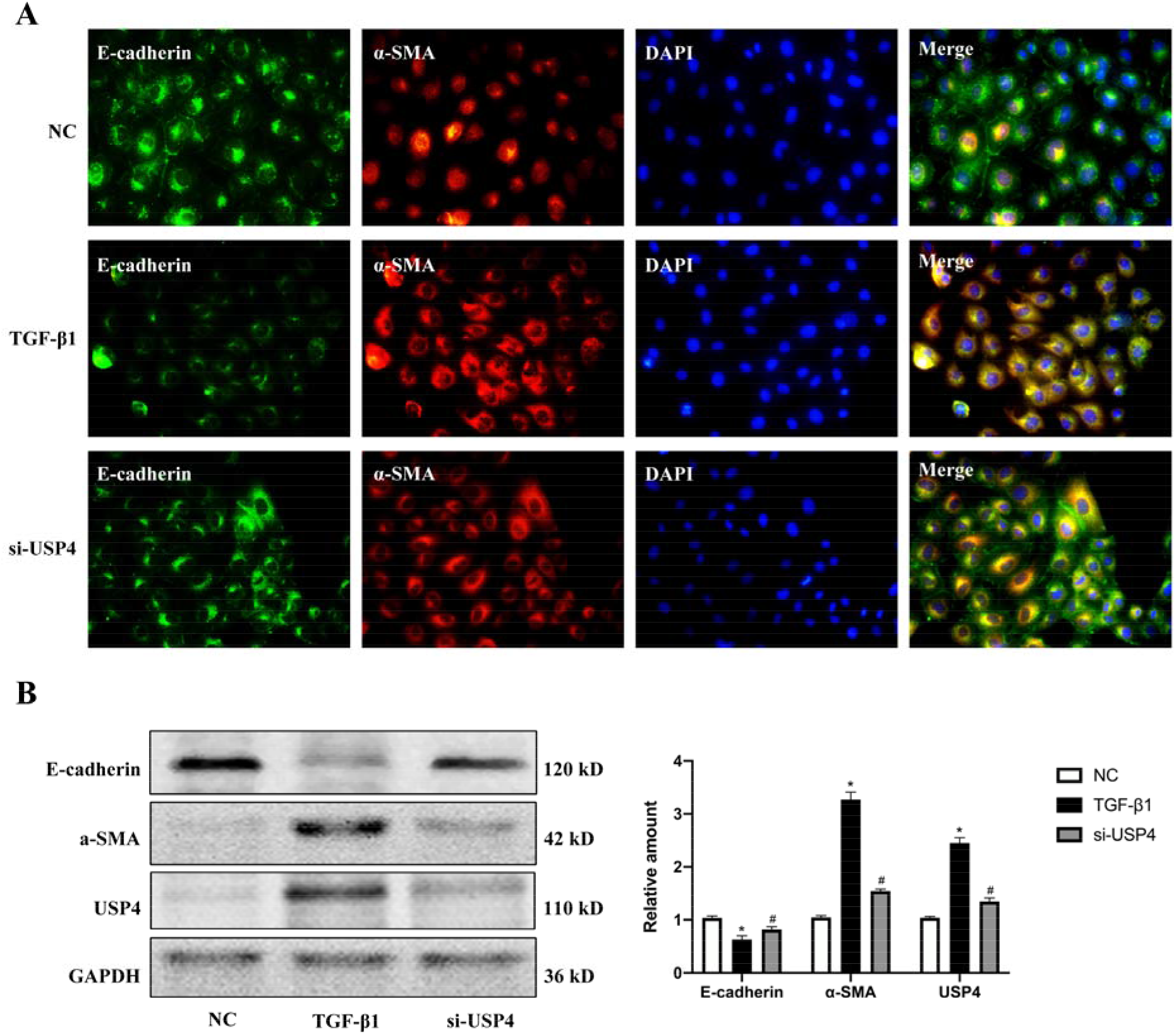
Effect of USP4 depletion on E-cadherin and α-SMA expression in NRK-52E cells. (A) Immunofluorescence microscopy showed downregulated E-cadherin and upregulated α-SMA expression in NRK-52E cells treated with TGF-β1 after 24h, and normalized with transfection with si-USP4 pretreatment. (B) Similar results showed in NRK-52E cells, as detected by Western blot analysis. * p<0.05 compared with normal control cells. # p<0.05 compared with TGF-β1-stimulated cells. NC: normal control, si-USP4: USP4 siRNA.

### 3.4 USP4 depletion downregulates TβRI expression induced by TGF-β1

In vitro, NRK-52E cells expressed TβRI rarely. Treated with TGF-β1 for 24h, NRK-52E cells overexpressed TβRI. Pretreated with si-USP4, TβRI was downregulated noticeably (Fig. 4A). Western blot analysis showed expression of TβRI was lower in NRK-52E cells transfected with si-USP4 than stimulated with TGF-β1 only (Fig. 4B). These results indicated that USP4 could stabilize TβRI in NRK-52E cells in response to TGF-β1.

**Fig. 4.**
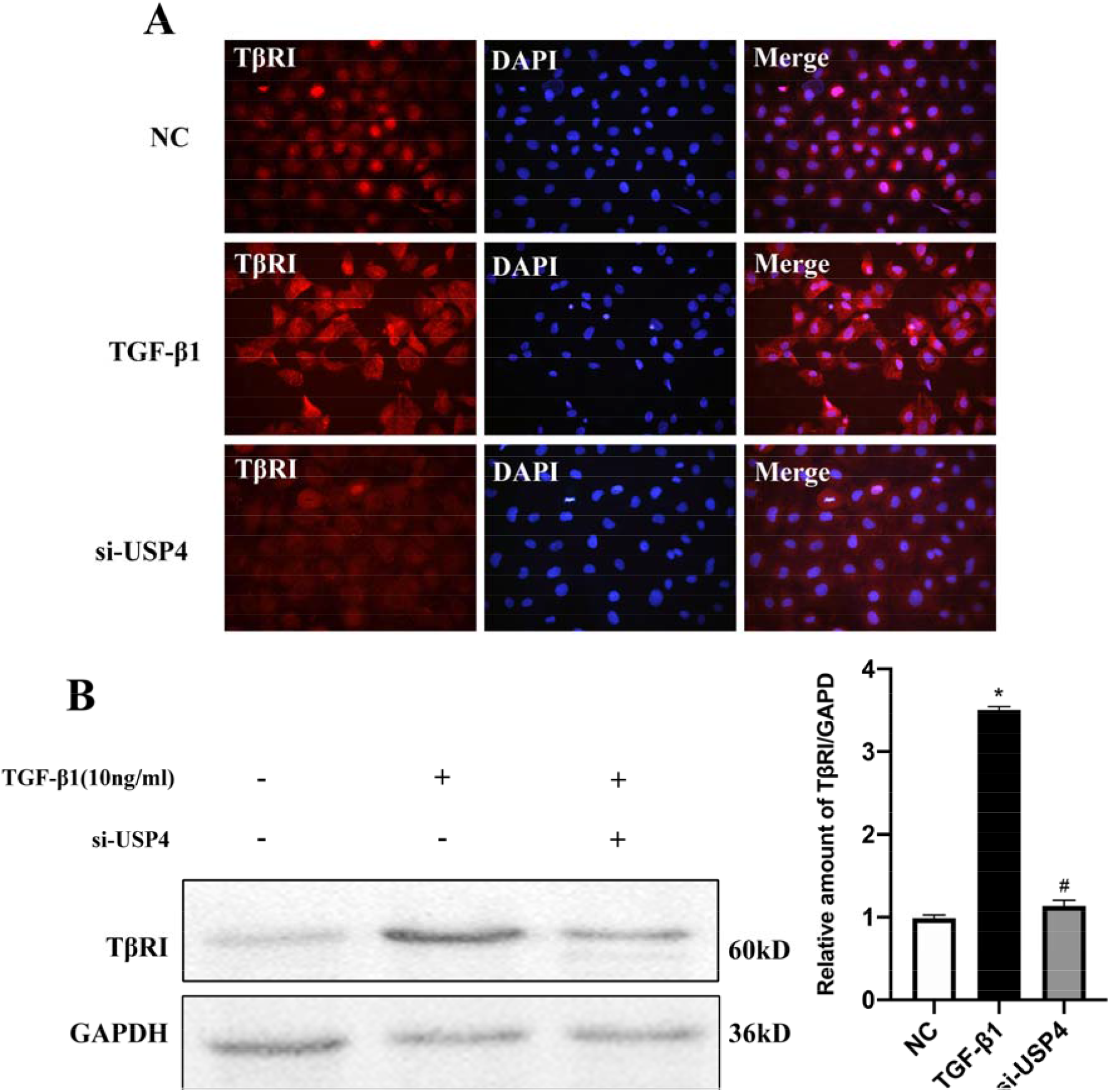
Effect of USP4 siRNA on TβRI expression in NRK-52E cells. (A) Immunofluorescence images of TβRI in NRK-52E cells as shown in images above, magnification ×400. (B) Western blot analysis of TβRI expression in NRK-52E cells, the bars show the relative expression quantity normalized to GAPDH and compared to the normal control group. * p<0.05 compared with normal control cells. # p<0.05 compared with TGF-β1-stimulated cells. NC: normal control, si-USP4: USP4 siRNA.

## 4. Discussion

This study shows that USP4 is an important molecule in the process of renal interstitial fibrosis. USP4 siRNA can decrease the expression of TβRI and improve the epithelial-mesenchymal transition of renal tubular epithelial cells. These results suggest that USP4 may be one of the important pro-fibrotic molecule in renal interstitial fibrosis.

Renal tubular epithelial cells are considered as the main target of various injuries and the initiator of fibrosis. Clinical studies have confirmed that the number of tubular epithelial cells with EMT characteristics is closely related to the severity of fibrosis and the level of serum creatinine in patients[17]. The prevention of EMT can delay the process of renal fibrosis. Our previous studies have also confirmed that EMT is closely related to renal interstitial fibrosis[18]. However, the cell lineage tracing method can not prove that epithelial cells can transform into myofibroblasts[4]. Recently, the concept of “partial EMT” is proposed. Partial EMT refers to the partial characteristics of mesenchymal cells acquired by epithelial cells, but not completely transformed into myofibroblasts[4]. In the process, the damaged tubular epithelial cells downregulate the epithelial marker protein and express the mesenchymal marker protein. Actually, it is believed that the tubular epithelial cells are not directly involved in renal interstitial fibrosis through the EMT process. Partial EMT indicates that although epithelial cells do not transform into myofibroblasts, their phenotype transformation is enough to affect the progress of fibrosis[19]. Therefore, inhibition of epithelial phenotype transformation can alleviate renal interstitial fibrosis.

The EMT process is activated by a variety of signaling pathways and regulated by epigenetic and post-translational modifications, such as methylation, acetylation, phosphorylation, glycosylation, hydroxylation and ubiquitination[4]. It is very important to study how post-translational modification plays a role in EMT regulation for discovering potential therapeutic targets of renal interstitial fibrosis. The function of the ubiquitin-protease system is closely related to many kidney diseases. The level of ENaC in epithelial cells is regulated by the ubiquitin-protease system, which is involved in the occurrence of Liddle syndrome[20]. The role of the ubiquitin-protease system has been confirmed in patients with chronic kidney diseases[21]. Trps1, a downstream effector molecule of BMP7, regulates E3 ubiquitin ligase Arkadia in renal interstitial fibrosis[22]. Smad ubiquitination regulator 2 degrades Smad2 and Smad7 through ubiquitin-dependent pathway, which plays a key role in regulating TGFβ/Smad signaling pathway[23]. Emerging evidences indicate that DUBs also participate in renal interstitial fibrosis[24][25][26]. Our study reveals that USP4 is associated with renal interstitial fibrosis. Otherwise, USP4 also participate in multiple organ fibrosis. It is reported that overexpression of USP4 can attenuate cardiac hypertrophy and fibrosis induced by angiotensin II and improve cardiac dysfunction in vivo by down-regulating the TAK/JNK/p38 signaling pathway[27]. USP4 inhibitor Vialinin A attenuates liver fibrosis through mTOR signaling pathway[28]. USP4 can promote scar formation after skin trauma[29]. The role of USP4 in cancer cells has been well studied. Recently, it has been found that USP4 mediates the deubiquitination of Twist1, an EMT-related transcription factor, to promote stem-like character of lung cancer cells[30]. USP4 activates hepatic stellate cells and EMT of hepatocytes[31]. USP4 also promotes mesothelial-mesenchymal transition induced by peritoneal dialysis by maintaining the stability of TβRI[32]. Some studies have confirmed that USP4 can promote proliferation, inhibit apoptosis, and facilitate chemoresistance of glioblastoma multiforme (GBM) cells[33][34]. Overexpression of USP4 mediates tumor growth and its correlation with malignant phenotype suggest that USP4 is a tumor-promoting factor in liver tumors[35]. We find that USP4 can regulate the EMT process of renal tubular epithelial cells. Considering its functional diversities of USP4, we presume that EMT is not only one of the mechanisms of regulating renal fibrosis. In this study, we discover that USP4 may play an important role in the EMT process of renal tubular epithelial cells during renal interstitial fibrosis.

The occurrence and development of EMT are affected by various factors during renal interstitial fibrosis. TGFβ1 is the most effective molecule in promoting renal interstitial fibrosis, and the activation of the TGFβ signaling pathway is considered to be the key initiating factor of EMT[36]. TGFβ1 exerts its role by binding with its downstream transmembrane receptor TβRI and TβRII to form activated complexes. TβRI is a key component of the TGFβ signaling pathway, located on the cell surface. After binding with TβRI, TGFβ1 affects the expression of downstream target genes through classical and nonclassical activation pathways[37]. It has been reported that the activation of TβRI can promote the expression of snail, twist, and other transcription factors, resulting in the decrease of E-cadherin expression in epithelial cells and the increase of mesenchymal markers, indicating an EMT[38]. Ubiquitination plays an important role in regulating and mediating TGFβ signal transduction[39]. TβRs are degraded by ubiquitination or lysosomal pathway. The stability and protein quantity of TβRI are strictly controlled by ubiquitination and mediate the activation of nonclassical signaling pathways[40]. TβRI is a ubiquitination target for degradation or cleavage, so it can downregulate TGFβ signal or stimulate the expression of target genes in the nucleus. Besides, activated TβRI recruits E3 ligase, which can initiate/regulate the signal transduction of TGFβ classical and nonclassical pathways. Different USPs regulate TGFβ pathway by stabilizing different factors in TGFβ pathway[39]. For example, USP4, USP11, and USP15 can stabilize the expression of TβRs by deubiquitination and enhance the epithelial-mesenchymal transition mediated by TGF-β1, thus leading to the metastasis of hepatocellular carcinoma, breast cancer, and glioblastoma respectively[41]. It is found that USP4 can directly bind to TβRI through its terminal, make its deubiquitination and upregulate the expression level of TβRI protein, thus enhancing the activity of TGFβ signaling pathway[13].

## 5. Conclusion

The results of this study confirm that USP4 interacts with TβRI in renal tubular epithelial cells. Inhibition of USP4 expression could increase the ubiquitination level of TβRI, increase the ubiquitin coupling of TβRI, thus increase the degradation of TβRI, reduce the expression of TβRI on the cell membrane and weaken the activity of the TGFβ signaling pathway. Downexpression of USP4 can decrease the EMT of renal tubular epithelial cells. The direct interaction between USP4 and TβRI promotes the EMT of renal tubular epithelial cells, which may be one of the cellular and molecular mechanisms for the development of renal interstitial fibrosis. The specific mechanism of USP4 to promote EMT of renal tubular epithelial cells needs further study.

