## Supplementary figures and images for "Inhibition of USP4 attenuates epithelial-mesenchymal transition of renal tubular epithelial cells by TβRI"

### Fig. 1.jpg

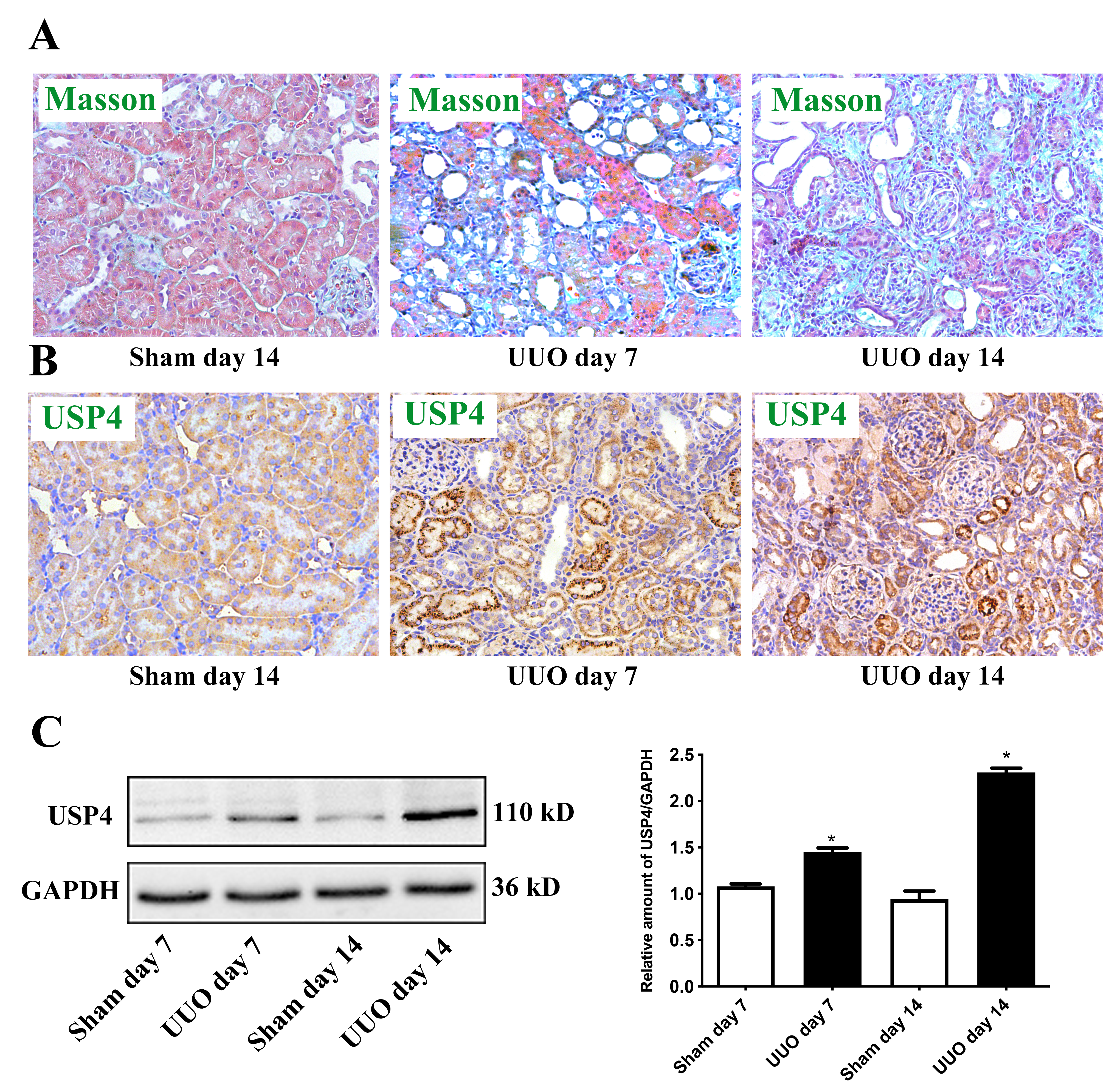

### Fig. 2.jpg

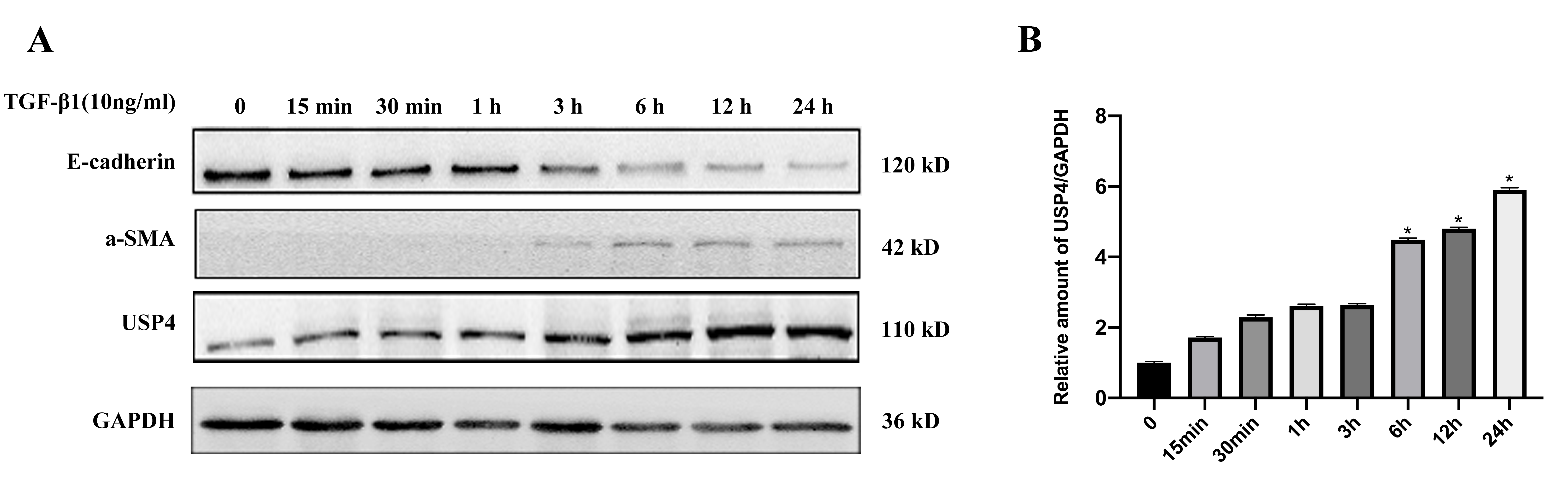

### Fig. 3.jpg

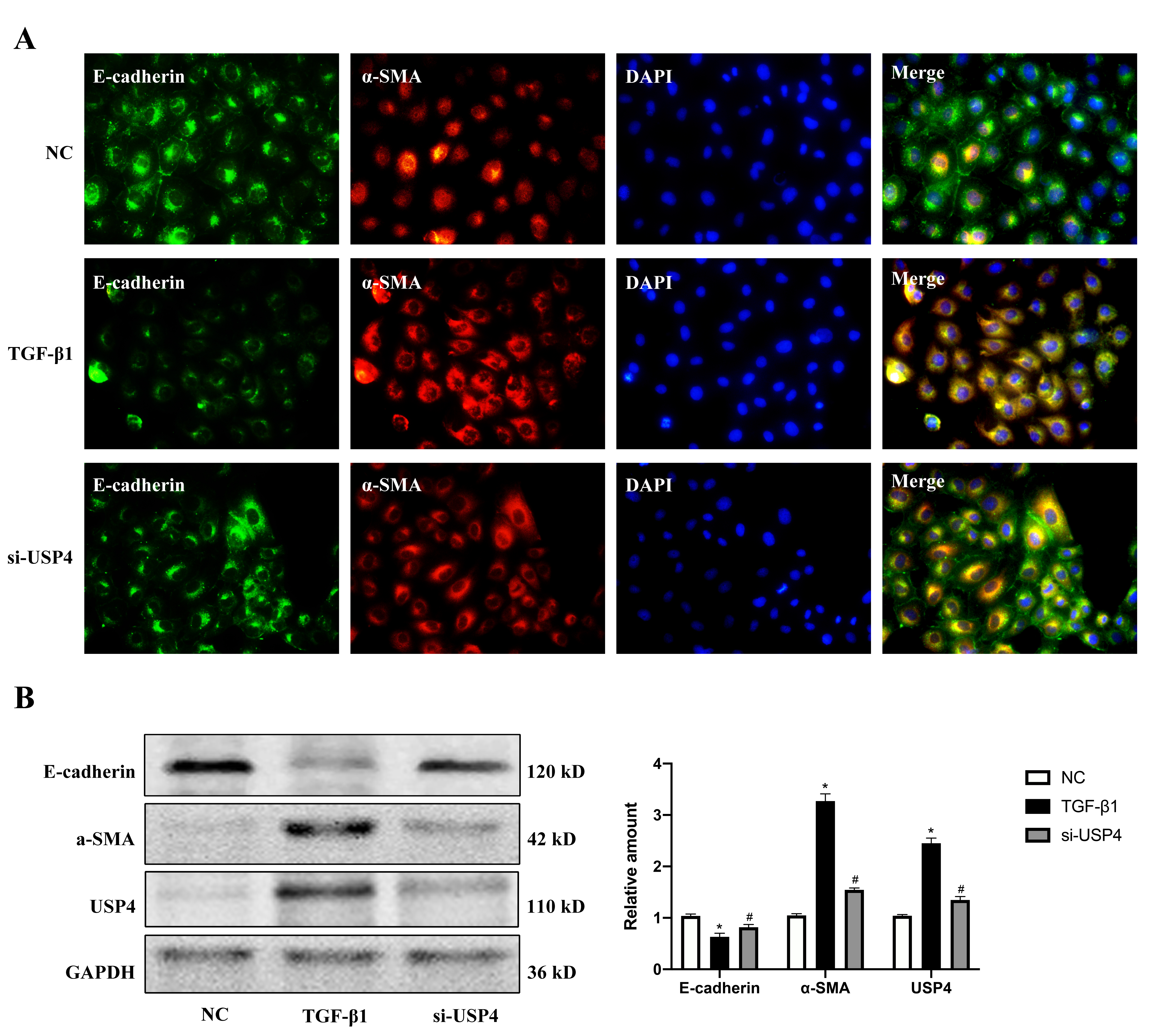

### Fig. 4.jpg

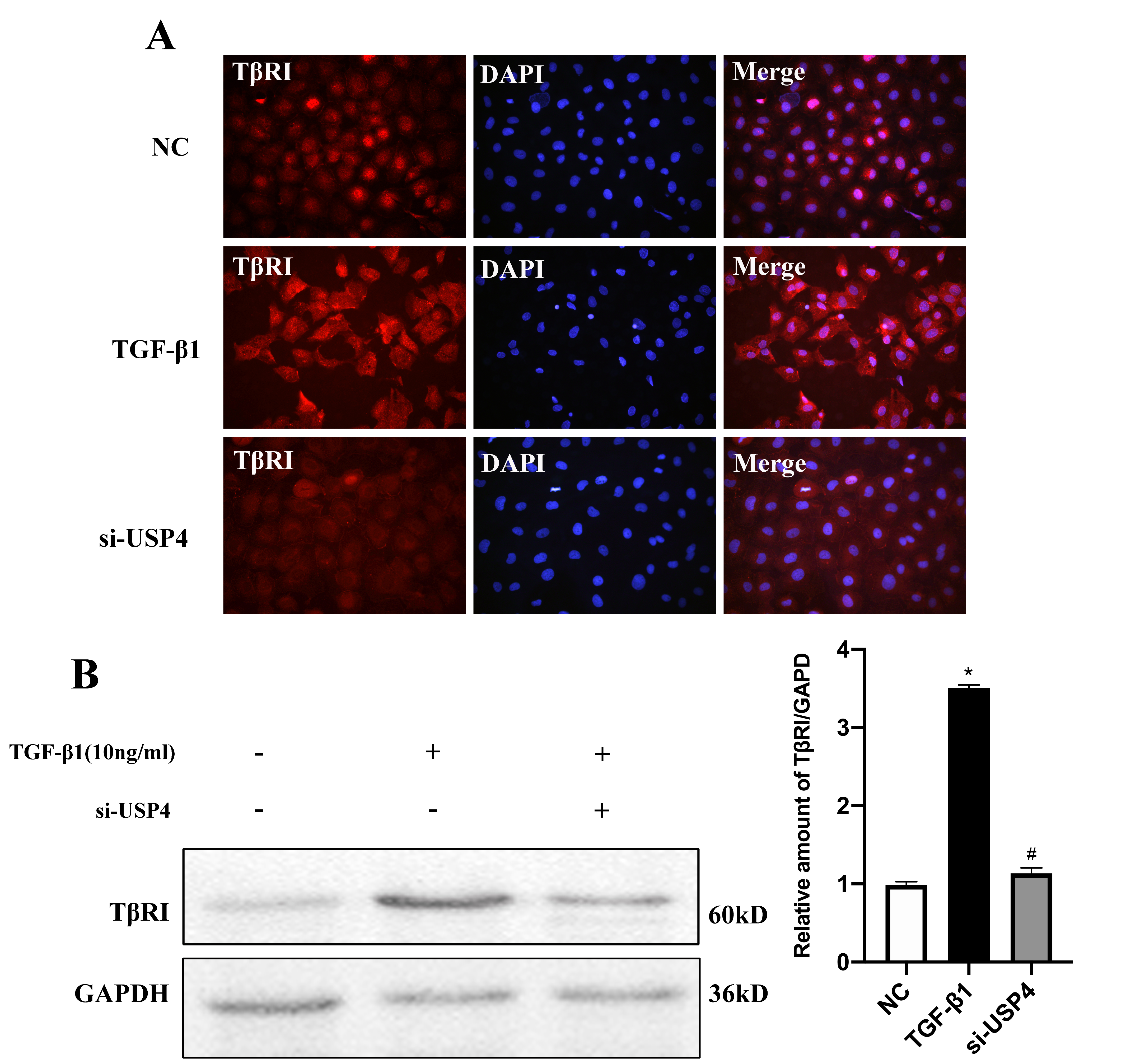
